# Neural representation of vocalization self-monitoring in the bat auditory midbrain

**DOI:** 10.1101/2024.09.12.612784

**Authors:** Huan Ye, Zhongdan Cui, Zhenxu Liu, Xuejiao Qin, Xincao Huang, Yuting Bai, Jinhong Luo

## Abstract

During acoustic interaction, mammals, including humans, not only monitor sounds from external sources but also their own vocalizations, the latter of which is known as vocalization feedback monitoring. Yet, it is unclear whether subcortical auditory regions process vocalization feedback. Here, we established an experimental paradigm that allows recording chronically the single-unit activity of the auditory midbrain (inferior colliculus) in freely vocalizing bats. We report that most collicular neurons represented self-produced biosonar vocalizations distinctively from pure tones and vocalization playbacks. Some neurons showed robust excitatory responses to vocalization feedback yet lacked any excitatory response to pure tones covering the entire frequency range of the vocalizations. Surprisingly, some collicular neurons even reversed the response polarity, from vocalization-induced suppression to vocalization-playback-induced excitation. Moreover, approximately a third of the neurons responded faster to self-produced vocalizations than to both vocalization playbacks and pure tones. These findings show that the midbrain inferior colliculus is involved in vocalization feedback processing, an ability suggested to arise locally in the auditory cortex.

## Introduction

Our knowledge of the mammalian auditory system is largely founded on the responses of auditory neurons to external sounds (i.e., exafference), in which the listening subjects were typically not vocalizing. During acoustic interaction, the auditory system not only processes external sounds generated by others but also the auditory feedback of self-produced vocalizations (i.e., reafference) ^1,2^. Auditory feedback monitoring plays a pivotal role in the real-time tuning of vocalization parameters, the longtime maintenance of vocalization structure, and in some species the acquisition of new vocalizations (i.e., vocal production learning) ^3,4^. However, there is only sparse research on the auditory neuronal activities in freely vocalizing mammals.

To date, non-human primates represent the dominant mammalian model from which the auditory neuronal activities have been recorded in freely vocalizing individuals ^5–8^. Consistent with observations in human subjects ^9,10^, these primate studies generally found that auditory cortical neurons exhibited suppressed activities during vocalization, i.e. the speaking- or vocalization-induced suppression. Studies on head-fixed, running mice also found that locomotion suppresses the auditory cortical, but not the thalamic, responses to movement-linked sound broadcasts and proposed that this suppression arose locally within the auditory cortex ^11,12^. Suppressed auditory responses to self-generated sounds are regarded as an adaptive mechanism to maintain the sensitivity of the brain to unpredictable external sounds during movement.

The IC is a central hub for auditory processing within the midbrain. It receives both ascending and descending inputs from most auditory nuclei ^13^. In addition, various non-auditory signals converge at the IC. These include signals related to vision, somatosensation, eye position, and behavioral context ^14^. Recently, investigations on rats have revealed that the primary motor cortex projects directly to the central nucleus of the IC ^15,16^. This raises the possibility that motor actions can directly modulate the auditory processing in the IC. At present, the single-unit activity of the IC neurons in freely vocalizing mammals is extremely limited. One study on the squirrel monkey reported that the neurons in the central nucleus of the IC did not discriminate self-produced from conspecific-produced vocalizations ^17^. Conversely, an earlier study on the greater horseshoe bat under anesthesia, in which the vocalizations were elicited by electrically stimulating the periaqueductal gray, found that some IC neurons responded differently to self-produced vocalizations compared to the playback stimuli that mimicked a portion of the intact vocalizations ^18^.

Here, we established an experimental paradigm to chronically record the single-unit activity of the IC neurons in freely vocalizing bats. Specifically, we recorded the IC activities in spontaneously vocalizing great roundleaf bat *Hipposideros armiger* hanging from a perch (**Figure 1A**). *H. armiger* is a highly vocal bat species that relies heavily on auditory feedback for online vocalization control ^19–22^. We report that, unlike the squirrel monkey, 88% of the IC neurons responded distinctively during the production of sonar vocalizations, compared to vocalization playbacks. The neural responses to vocalization feedback and vocalization playbacks differed not only quantitatively in 52% of the neurons, but also qualitatively in 43% of the neurons.

**Figure 1.**
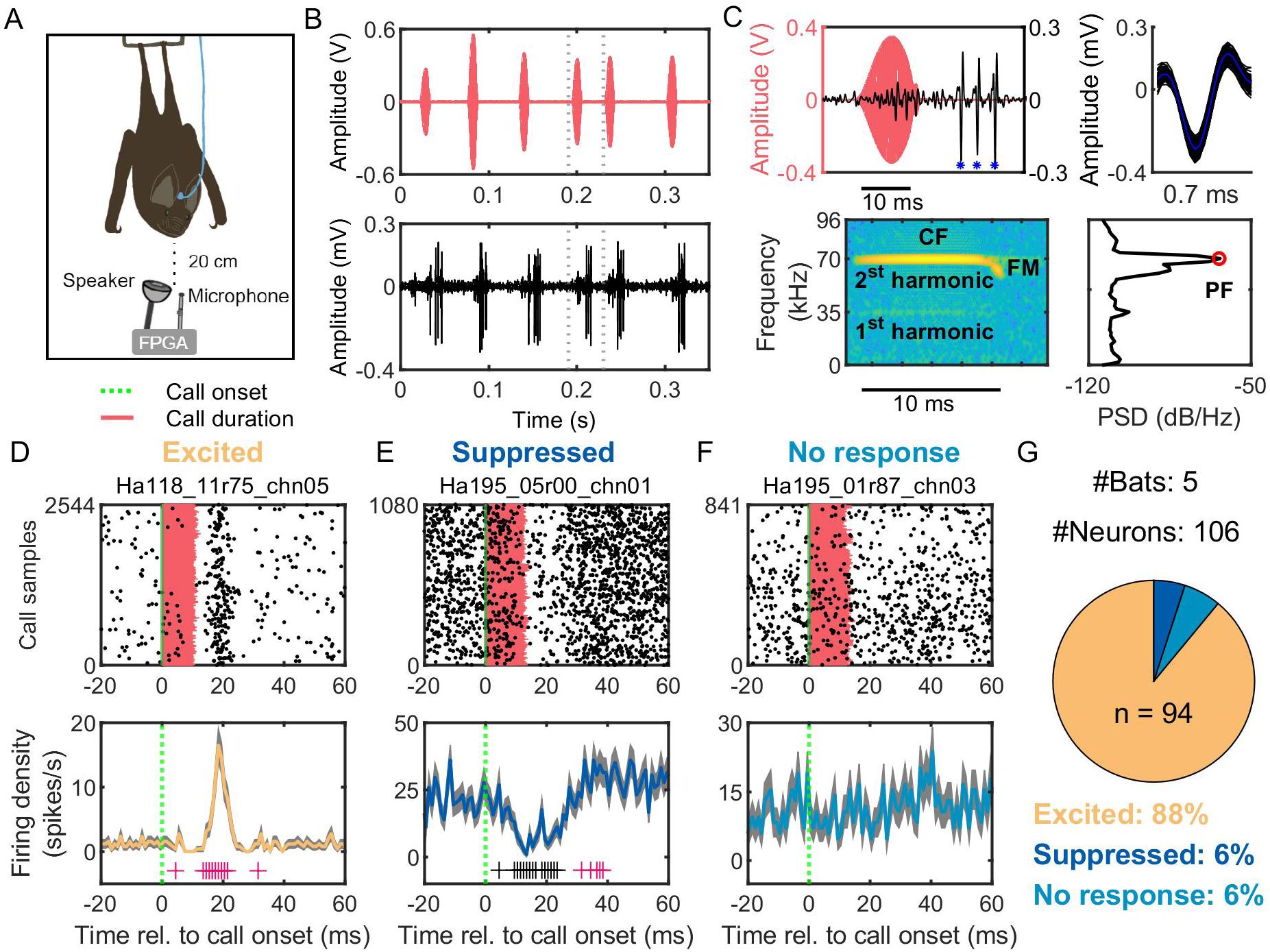
An experimental paradigm for studying the neuronal activity in freely vocalizing bats. (A) An illustration of the recording setup. A bat, *Hipposideros armiger*, hung on a perch and emitted sonar vocalizations that were recorded by a calibration microphone. The neuronal activity was recorded chronically with an implanted 16-channel silicon probe. The sound stimulus can be played back to the bat via a loudspeaker at a distance of 20 cm. (B) An example of the synchronized recordings of the self-emitted sonar vocalizations and the neuronal activity in the inferior colliculus (IC) in *H. armiger*. (C) A zoom-in view of the IC spike activity (blue asterisks) associated with a sonar vocalization (red). The spike shape is shown in the right panel. The spectrogram and power spectrum of this vocalization are shown in the bottom panel. The red circle was the peak frequency (PF) of this vocalization. (D, E, F) Raster (upper panels) and PSTH (1 ms bin size) plots (bottom panels) of an excitatory neuron (D), a suppressive neuron (E), and a no-response neuron (F) of the bat IC. Error bars indicate SEM (shaded). Bins with significant change from baseline are indicated (carmine represents time bins with significant increases; black represents significant decreases; *p* < 0.05, rank-sum test with FDR correction). (G) Most neurons showed excitatory responses to self-emitted sonar vocalizations.

## Results

We recorded the neuronal activities of the IC using implanted 16-channel silicon probes from freely vocalizing *H. armiger*. During a recording session, each *H. armiger* spontaneously produced thousands of biosonar vocalizations (echolocation calls) while hanging at a perch (**Figure 1A**). The biosonar vocalizations were recorded by a nearby ultrasonic microphone synchronously with the neural activities (**Figure 1BC**). In addition, acoustic stimuli can be broadcasted to the bat via a calibrated ultrasonic speaker. In total, we isolated 106 single units from five implanted bats (**Figure 1C**, upper panel, see **Table S1** for details). The sonar vocalizations of *H. armiger* were short (∼10 ms), and typically comprised of a constant-frequency (CF) component followed by a downward frequency-modulated (FM) component (**Figure 1C**, lower panel). Most energy of the sonar vocalizations in *H. armiger* concentrates in the 2^nd^ harmonic and the peak frequency is around 70 kHz.

One of the first observations, that was already evident during the experimental recording sessions, was that many IC neurons exhibited strong excitatory responses to self-produced sonar vocalizations. The neural activities seemed to be phasically locked to each sonar call emission (**Figure 1B**) and may appear within a few milliseconds after the onset of the sonar vocalization (**Figure 1C**; the small unsorted units during the call production). When aligning the neural activities to the onset of sonar vocalizations, we observed that many neurons fired strongly after the onset of the sonar vocalizations within a restricted time window (**Figure 1D**). A few neurons decreased the neural activities to self-produced vocalizations (**Figure 1E**), and some did not change the activities after self-produced vocalizations (**Figure 1F**). Overall, 88% (94/106) of the neurons were excited, 6% (6/106) were suppressed, and 6% (6/106) were not affected by self-produced vocalizations (**Figure 1G**).

To understand whether the collicular responses to self-produced vocalizations can be explained as pure auditory responses to acoustic stimuli, we characterized the frequency tuning curves of most recorded IC neurons (n = 97). 91 of these neurons were characterized by both pure tones and self-produced vocalizations. The pure tones matched the main acoustic features of the echolocation calls: they were 10 ms in duration, covered the frequency range from 5 kHz to 90 kHz at a step of 1 kHz, and were played at 75 dB SPL peak amplitude. This playback amplitude is probably comparable to or even slightly greater than the perceived amplitude of the self-produced calls (see **Methods**). While most neurons showed excitatory responses to pure tones at selective frequencies (**Figure 2A**), some neurons did not respond to pure tones at all (**Figure 2B**). From the excitatory frequency tuning curves, we measured the best frequency and response peaks. Best frequency refers to the pure tone frequency that evoked the highest excitatory response (marked by the unfilled triangle in **Figure 2A**). Some neurons contained multiple response peaks (marked by unfilled circles in **Figure 2A**) across the tested frequencies of pure tones and these neurons were known as multi-peak neurons ^23,24^. Overall, 92% (89/97) of the neurons contained excitatory response peaks and thus best frequency to pure tones (**Figure 2A**). When mapping the best frequency and response peaks to the frequency range of the sonar vocalizations of *H. armiger*, we found that the best frequency in 78% of the neurons fell within the frequency range of the 2^nd^ harmonic of the sonar vocalizations, with an additional 10% of neurons in the frequency range of the 1^st^ harmonic (**Figure 2C**). However, 39% of these neurons also contained response peaks beyond the spectral ranges of the sonar vocalizations (i.e., the 1^st^ and the 2^nd^ harmonics) (**Figure 2D**).

**Figure 2.**
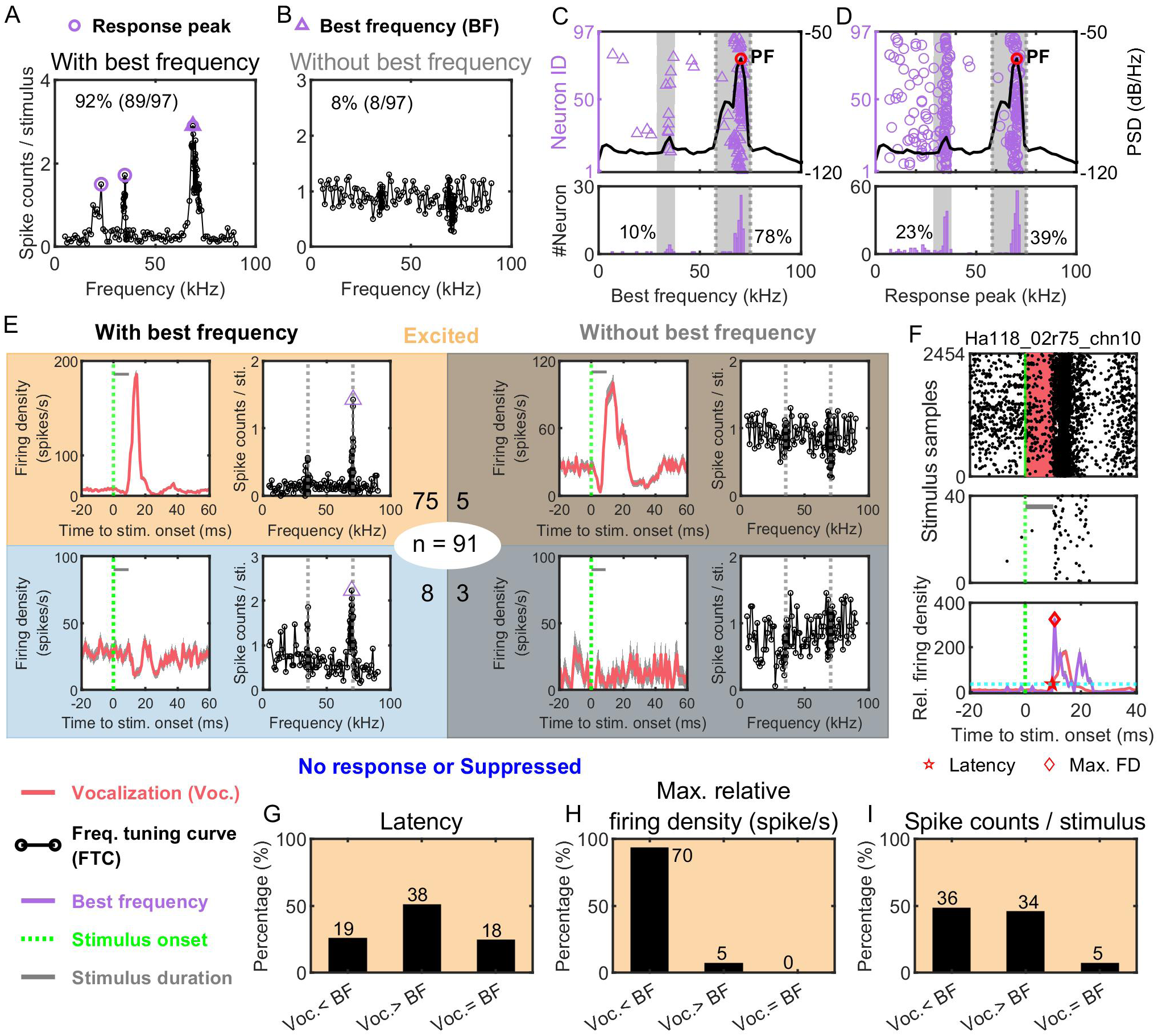
Most inferior colliculus (IC) neurons exhibited strong excitatory responses to pure tone. (A) An example neuron showing maximum excitatory responses to pure tones at a given frequency, which was referred to as the best frequency (BF). Unfilled triangles indicate the best frequency, and circles indicate the response peaks. Numbers in the upper left corner indicate the proportion of neurons of each type. (B) An example neuron that did not respond to pure tones, and thus lacked the best frequency. (C) The distribution of the best frequency with the frequency ranges of the sonar vocalizations. The bottom panel indicates the percentages of frequency distributions within the frequency range of the first and the second harmonics. (D) The distribution of the response peaks with the frequency ranges of the sonar vocalizations. (E) Four types of neurons are classified by qualitatively comparing the responses to self-produced vocalizations and pure tones. 89 neurons (in the ellipse) were examined by both the self-produced vocalizations and pure tones. Numbers surrounding the ellipse indicate the number of neurons of each neuron type. The median duration of the vocalizations is indicated by the length of the gray bar. (F) Raster (upper and middle panels) and PSTH (1 ms bin size) plots (bottom panels) of an example neuron in response to self-produced vocalizations and the pure tone at the best frequency. Star and diamond indicated the latency and maximum firing rate at the best frequency, respectively. (G-I) Pairwise comparisons of the latency, the maximal relative firing density, and the spike counts per stimulus of the IC neurons showed excitatory responses to both self-emitted vocalizations and pure tone at the best frequency (as in the top left panel of **E**). The three response parameters were measured by randomly drawing 100 calls from the entire vocalization dataset, or 30 pure tones from the entire pure tone dataset. For each neuron, we repeated this process 30 times and evaluated the statistical significance of the responses between self-produced vocalizations and pure tones at the best frequency (rank-sum test, *p* < 0.05).

By qualitatively comparing the responses evoked by self-produced vocalizations to those by pure tones, we classified the neurons into four types (**Figure 2E**). Type I neurons (n = 75), which is the most common type, showed excitatory responses to both self-produced vocalizations and pure tones. Type II neurons (n = 5) showed strong excitatory responses to self-produced vocalizations but did not respond to pure tones. Type III neurons (n = 8) showed suppressed responses to self-produced vocalizations, but excitatory responses to pure tones. Type IV neurons (n = 3) did not respond to either self-produced vocalizations or pure tones and thus represented non-auditory, spontaneously firing neurons in the present study.

Next, we tested whether the responses to self-produced vocalizations and pure tones at the best frequency can be differentiated for the Type I neurons, despite the similarity in the gross firing pattern. For each neuron, we measured two response parameters: the response latency and the maximum relative firing density (relative to the background firing density in the 20 ms window before stimulus onset) (**Figure 2F**). We found that 24% (18/75) of the neurons had statistically indistinguishable response latency to self-produced vocalizations and the pure tone at the best frequency (**Figure 2G**). By contrast, 93% (70/75) of the neurons had higher maximum relative firing densities to the pure tone at the best frequency than self-produced vocalizations (**Figure 2H**). However, the proportion of neurons that showed a higher firing rate (spike counts per stimulus) to self-produced vocalizations than pure tones was similar to that showed a lower firing rate (**Figure 2I**). This result suggests that the much higher proportion of neurons that had higher firing density to pure tones can be attributed to the invariable parameter space of pure tones that resulted in reduced jittering in response latency.

To understand whether the above response differences in the IC neurons between self-produced vocalizations and pure tones reflect purely acoustic differences between these two groups of stimuli, which may explain the result in the greater horseshoe bat ^18^, or the motor action linked to self-produced vocalizations, we compared the responses of some IC neurons to self-produced vocalizations and vocalization playbacks when the bat was not actively vocalizing, i.e. when the bat was in a relatively quiet behavioral state. The vocalizations used for playback were from the same bat and contained both the 1^st^ and 2^nd^ harmonics (**Figure 3A**; see Methods for details). A total of 40 single-units were recorded from two bats and 88% of these neurons were excitatory to self-produced vocalizations (**Figure 3B**). Overall, we found that 43% of the neurons exhibited distinct response patterns to self-produced vocalizations and vocalization playbacks in a qualitative way (**Figure 3C-F, J**). Qualitative differences in neuronal response pattern included a change from vocalizations-induced single excitatory peak to playback-induced two excitatory peaks (**Figure 3C; 17%**), a change from vocalizations-evoked two excitatory peaks to playback-induced single excitatory peak (**Figure 3D; 8%**), a change from vocalizations-evoked suppressive response to playback-evoked excitatory response (**Figure 3E; 10%; Movie S1**), and a change from vocalizations-evoked excitatory response to playback-evoked suppressive or no response (**Figure 3F; 8%**). The recording depths of the neurons that reversed the response polarity were between 1, 882 µm and 3, 454 µm, which correspond to the central portion of the IC (**Figure S1**). 5% of the neurons did not show responses to either self-produced vocalizations or vocalization playbacks (**Figure 3I**).

**Figure 3.**
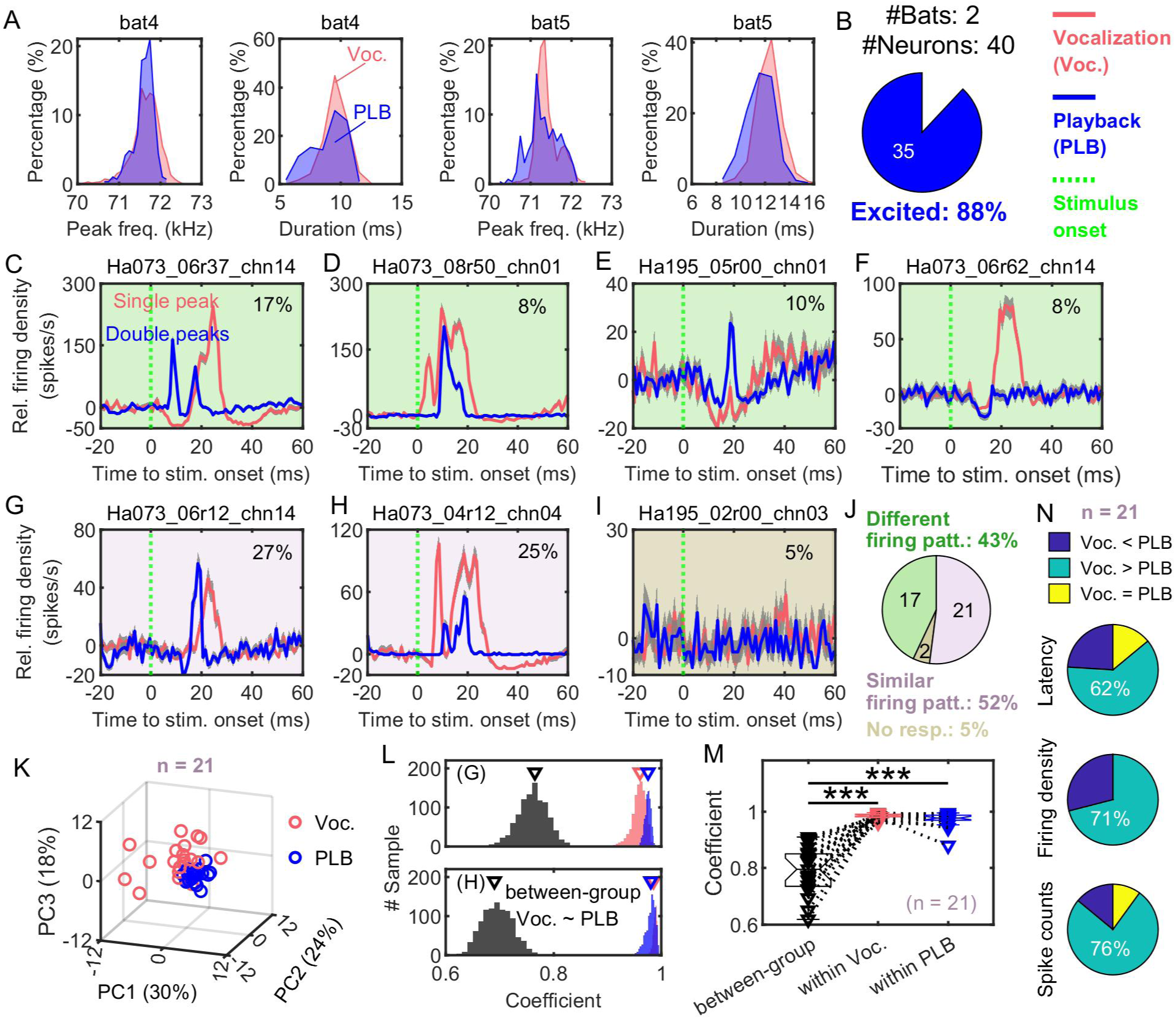
Most inferior colliculus (IC) neurons represented self-produced vocalizations distinctively from vocalization playbacks. (A) The distribution of peak frequency and duration of self-produced vocalizations and the vocalizations used for playback were selected from the same bat. (B) Most neurons showed excitatory responses to vocalization playback. (C-I) Distinct response patterns of the neurons to self-produced vocalizations and vocalization playbacks. Numbers in the upper right corner indicate the proportion of neurons of each type. (C, D) Two example neurons show the qualitative changes of response patterns between double-peaks and single-peak. (E, F) Two example neurons show the reversed response polarity between suppressive and excitatory. The recording depths were 2990 µm and 2882 µm, respectively. (G, H) Neurons that showed qualitatively similar firing patterns to self-produced vocalizations and vocalization playbacks (n = 21 neurons). (I) The neurons did not show responses to either self-produced vocalizations or vocalization playbacks, which were referred to as the no-response group. (J) Proportions of different firing pattern groups in (C-I). firing patt., firing pattern, No resp., no response. (K-N) Quantitative analysis for neurons showing similar firing patterns in (G) and (H). (K) Scatter plots of the spike data (40 ms PSTH after the onset of stimulus) for vocalizations or vocalization playbacks using the first three principal components (PCs), based on data in (G) and (H). (L) Distributions of the coefficients from the cross-correlation analysis for self-produced vocalizations and vocalization playback, and between vocalizations and vocalization playback for two example neurons (0.005 bin size, the top panel was for data shown in (G), and the bottom panel was for data shown in (H)). The triangles indicate the peak of the distributions. (M) The comparison of median coefficients for all neurons between the experimental groups. (rank-sum test, ***, *p* < 0.001). (N) Proportions of neurons based on statistical comparisons of the latency, the maximal relative firing density, and the spike counts per stimulus of IC neurons in response to self-produced vocalizations and vocalization playbacks. The three response parameters were measured by resampling to the 100 randomly selected calls and the 100 randomly selected vocalization playbacks from the entire vocalization and vocalization playbacks dataset respectively. For each neuron, we repeated this process 30 times and evaluated the statistical significance of the responses between self-produced vocalizations to vocalization playbacks (rank-sum test, *p* < 0.05).

Then, we tested whether the neurons that showed qualitatively similar firing patterns to self-produced vocalizations and vocalization playbacks (**Figure 3GH**; 52%) could be distinguished quantitively. We performed the principal component analysis (PCA) using the spike data (40 ms PSTH after the onset of vocalizations or vocalization playbacks). We found that the neuronal responses to self-produced vocalizations and vocalization playbacks can be differentiated for 72% of the units already by the first three principal components (**Figure 3K**). As this PCA analysis used the absolute firing pattern, two firing patterns will be differentiated when either the firing rate or response latency is different, despite the similarity in the general firing profile. For many IC neurons, a change in stimulation amplitude can modify the firing rate and/or response latency, yet during the experiment, it is virtually impossible to ensure that the perceived vocalization amplitude would perfectly match the vocalization and playback conditions. To understand to what extent the elicited neuronal differences by self-produced vocalizations and vocalization playbacks may be simply due to a difference in stimulus amplitude, we further used the normalized cross-correlation method to compare similarity in firing pattern. Specifically, we performed the similarity analysis with 50% of the vocalizations and 50% of the playback vocalizations randomly sampled from the experimental datasets. Then we calculated the maximum correlation coefficient for the firing patterns between the resampled 50% vocalization dataset and original vocalization dataset (within-group analysis), between the resampled 50% playback vocalization dataset and the original playback dataset (within-group analysis), and between the resampled 50% vocalization dataset and the resampled 50% playback vocalization dataset (between-group analysis). This analysis was repeated 1000 times. We found that the peak correlation coefficients, and thus the response similarity, were significantly smaller between the stimulation groups than within the same group (**Figure 3LM**). Furthermore, 62% (13/21) of these neurons had statistically longer response latency to self-produced vocalizations than vocalization playbacks (**Figure 3N**). 71% (15/21) and 76% (16/21) of these neurons had higher maximum relative firing densities and spike counts to self-produced vocalizations than vocalization playbacks. Thus, the differences in firing patterns between the vocalization and vocalization playback conditions were not solely due to differences in the maximum firing rate or response latency that may be associated with differences in the stimulation amplitude.

Lastly, we compared the neuronal responses in a portion of neurons (n = 22) that were excited by all three stimulus types (22/37, 59%), i.e. self-produced vocalizations, vocalization playback, and pure tones. The responses of one example neuron is shown in **Figure 4A**. This neuron had an approximately 6.7 and 4.8 ms shorter response latency and a broader response window to self-produced vocalizations than to vocalization playback and the pure tone at the best frequency (squares marked in **Figure 4B**). Overall, we found that 32% (7/22) of the IC neurons had a shorter response latency to self-produced vocalizations than to both vocalization playback and pure tone at the best frequency statistically (**Figure 4B**). The median response latency of these neurons to self-produced vocalizations was 3.0 ms shorter (ranging from 0.7 to 6.7 ms) than to vocalization playback and was 1.8 ms shorter (ranging from 0.7 to 4.7 ms) than to the pure tone at the best frequency. Interestingly, almost all neurons (8/9) with shorter latency to self-produced vocalizations contained two excitatory peaks. Compared to the uniformly shorter response latency in the first excitatory peak, the response latency of the second excitatory peak of these two-peak neurons was either shortened, lengthened, or not changed. Furthermore, 64% (14/22) of the IC neurons responded with a higher firing rate to self-produced vocalizations not only than to vocalization playback but also to the pure tone at the best frequency statistically (**Figure 4C**). Based on the median, for each stimulus, these neurons fired 1.1 spikes more to self-produced vocalizations than to vocalization playback and 0.8 spikes to the pure tone at the best frequency. Furthermore, we found that there was a strong negative correlation between the response latency and firing rate in these neurons in response to only self-produced vocalizations (**Figure 4D**, pink circles in the left panel; Pearson Correlation, *r* = -0.67, *p* < 0.001), but not in response to either vocalization playback (**Figure 4D**, magenta circles in the left panel; Pearson Correlation, *r* = -0.05, *p* = 0.84) or pure tone at the best frequency (**Figure 4D**, blue circles in the left panel; Pearson Correlation, *r* = -0.22, *p* = 0.31). This unique negative correlation between response latency and firing rate in the self-produced vocalization condition was also evident with the entire sampled population of IC neurons (**Figure 4E**, pink circles in the right panel, Pearson Correlation, *r* = -0.41, *p* < 0.001).

**Figure 4.**
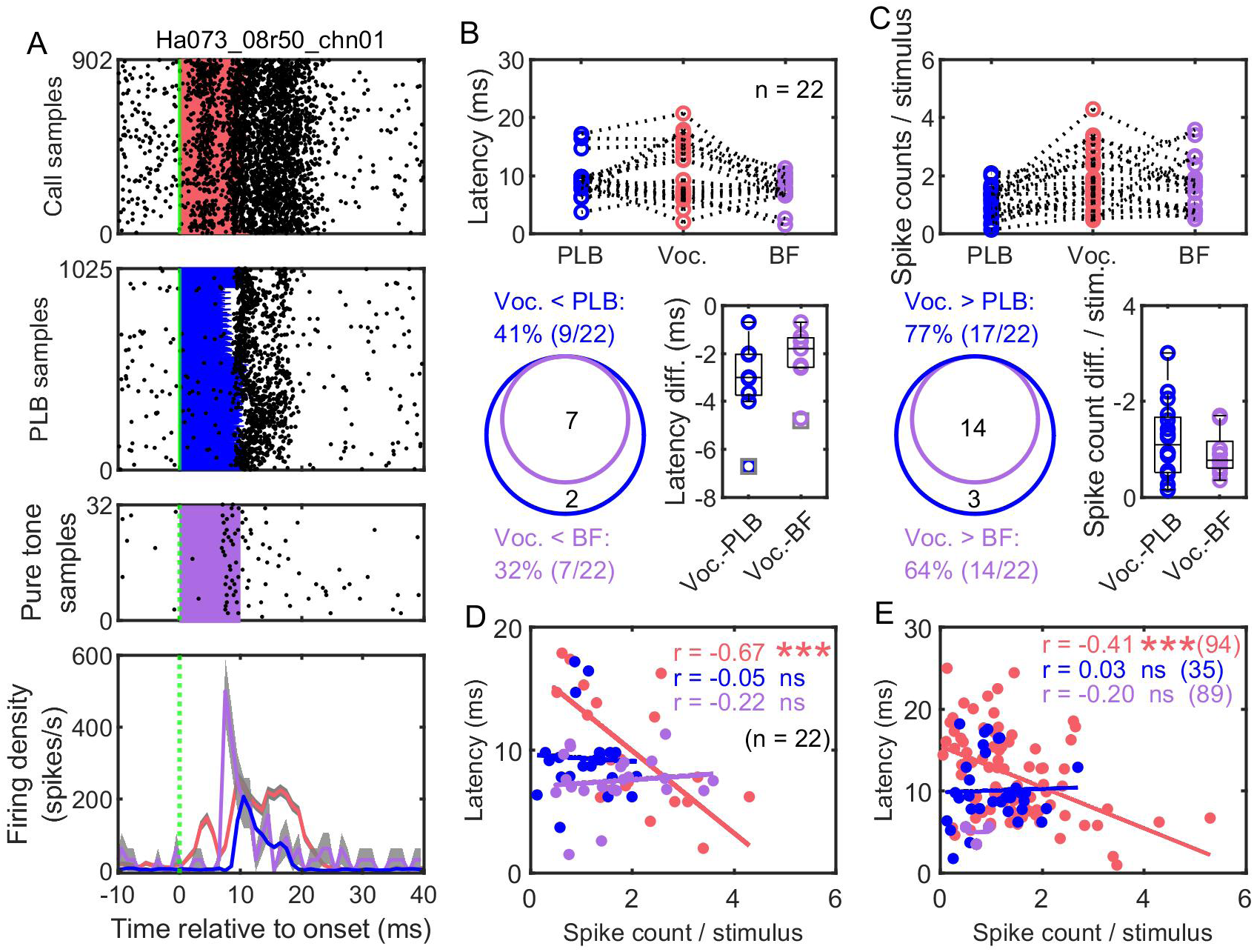
Inferior colliculus neurons exhibited reduced response latency to self-produced vocalizations than to vocalization playback and pure tones. (A) Raster (upper three panels) and PSTH (1 ms bin size) plots (bottom panel) of an example neuron in response to self-produced vocalizations, vocalization playback, and pure tone at the best frequency. (B) Comparison of the response latency in neurons (n = 22) that were excited in all three stimulus types (22/37, 59%). (C) Comparison of the spike count per stimulus in neurons that were excited in all three stimulus types. (D) Correlation between the response latency and spike count per stimulus in neurons in response to self-produced vocalizations, vocalization playbacks, and pure tone at best frequency (n = 22). (E) Correlation between the response latency and spike count per stimulus for all neurons in response to self-produced vocalizations (n = 94), vocalization playbacks (n = 35), and pure tone at best frequency (n = 89) (Pearson Correlation, ***, *p* < 0.001, ns; *p* > 0.05).

## Discussion

Does our auditory system function in a similar way when it processes sounds originating from external sources (e.g., conspecifics) and sounds originating from ourselves? Here we provide evidence that even at the level of the midbrain many IC neurons in a mammal, the echolocating bat *H. armiger*, responded dramatically differently to self-produced sonar vocalizations and playbacks of its own vocalizations. The differences in neural activities not only manifested as quantitative changes in the response profile of the same pattern, i.e. the firing density over time but also as reversed response polarity in nearly half of the IC neurons (43%, **Figure 3J**), such as from suppressive to excitatory. Furthermore, one important finding from the present study is that a relatively large population of the IC neurons (41%) responded more rapidly to self-produced vocalizations than vocalization playback and pure tones at the best frequency (**Figure 4B**). These results show that vocalization production exerts a strong influence on the auditory processing already in the midbrain inferior colliculus.

Our results contrast with the study on the squirrel monkey which found little difference in the neuronal responses within the central nucleus of the IC to self-produced vocalizations and vocalizations from loudspeakers or conspecifics ^17,25^. This difference is not due to the possibility that our recorded IC neurons were not from the central nucleus of the IC. In all our electrode implants, we have targeted the central portion of the IC which is visible from the skull for *H. armiger* (**Figure S1**). Due to the limited research to be compared, it is impossible to tell whether our observations were specific to echolocating bats. The IC of echolocating bats is hypertrophied in size and plays an essential role in echolocation ^26^. Critical functions of echolocation, including the echo-delay tuned neurons implicated for target ranging, emerge first in the IC along the ascending auditory pathway ^27–29^. Furthermore, echolocating bats routinely perform remarkably rapid audio-vocal control at a time scale within 100 ms or so ^30–33^, suggesting that essential neural circuits for echolocation probably lie in the midbrain and brainstem. These considerations might have strengthened the audio-vocal interaction in the IC of echolocating bats.

At present, the precise mechanisms for the difference in response to self-produced vocalizations and vocalization playbacks are unknown. We propose two mutually non-exclusive hypotheses: the efference copy/corollary discharge hypothesis and the behavioral state hypothesis. In the efference copy hypothesis, we suggest that vocal motor signals are sent to the IC shortly before or during active vocalization and these motor signals modulate the neuronal responses in the IC. Evidence supporting this hypothesis was from recent investigations on rats, which found that the primary motor cortex projects directly to the central nucleus of the IC ^15,16^. Consistent with the rat research, another study on head-fixed mice found that neural activities of IC neurons were strongly modulated by locomotion ^34^. Furthermore, our data revealed that a small percentage, up to 10%, of the IC neurons exhibited suppressed responses during self-produced vocalizations, but excitatory responses to playbacks of pure tones (**Figure 2G**) or vocalizations (**Figure 3E**). This class of neurons resembles the well-known vocal-feedback monitoring neurons in the auditory cortex of primates and mice, although the percentage of neurons showing vocalization-induced suppression in the auditory cortex is several times higher ^6,7,35,36^. It was argued that vocalization-induced suppression in the auditory pathway might be an indication of the corollary discharge signals originating from the vocal-motor pathway ^37^. These data thus challenge the hypothesis that the ability to suppress the predictable auditory feedback originates locally within the auditory cortex ^1,11^.

In the behavioral state hypothesis, we suggest that vocalization represents an active behavioral state distinct from that when the animal is not vocalizing. Emerging evidence showed that non-auditory signals, including task engagement, behavioral state, and attention, modulate the neural activity of the IC ^38–41^. The difference between active vocalization and passive listening can be particularly striking for echolocating bats ^42^. During active sonar vocalizing, echolocating bats probe and acquire information from the surroundings on purpose. During passive listening, like in other non-echolocating animals, the behavioral state in the listening subjects can vary substantially from active engagement to unconscious drowsiness. Future circuit level research using model species such as laboratory mice can be essential to test the causal relationship between vocal motor activites and subcortical auditory processing.

## Acknowledgments

We thank Aoqiang Li for help with bat catching, and Songlin Huang and Huanhuan Li for help with the setup. We thank professor Lixia Gao for valuable suggestions. This study was funded by a Career Development Award from the Human Frontier Science Program (CDA00009/2019-C), the National Natural Science Foundation of China (31970426; 32270535), and the Fundamental Research Funds for the Central Universities (CCNU22QN009; CCNU22LJ003) to JL.

## Author Contributions

HY, ZC, ZL, XQ, XH, and YB conducted the experiments; HY, ZC, and ZL analyzed the data; JL and HY wrote the manuscript; JL conceived, designed, and supervised the experiment.

## STAR Methods

### KEY RESOURCES TABLE

| REAGENT or RESOURCE | SOURCE | IDENTIFIER |
| --- | --- | --- |
| Experimental models: Organisms/strains |  |  |
| *Hipposideros armiger* | A cave in Xianning County, Hubei Province, China. | NA |
| Software and algorithms |  |  |
| MATLAB R2019b | MathWorks | RRID: SCR_001622 |
| LabVIEW | National Instruments | https://www.ni.com/zh-cn.html |
| NeuroStudio | Greathink Medical Technology | http://www.greathink.com/ |
| Neural data analysis scripts | This paper | DOI: 10.17632/vy4s3kyc97.1 |
| SoundMexPro toolbox | Hörzentrum Oldenburg | https://soundmexpro.hz-ol.de/index.php?page=3 |

## RESOURCE AVAILABILITY

### Lead contact

Further information and requests for resources and reagents should be directed to and will be fulfilled by the lead contact, Jinhong Luo

### Materials availability

This study did not generate new unique reagents.

### Data and code availability

The datasets supporting the current study have not been deposited in a public repository because of their large size, but are available from the corresponding author on request.

The original code has been deposited at Mendeley and is publicly available as of the date of publication. DOIs are listed in the key resources table.

Any additional information required to reanalyze the data reported in this paper is available from the lead contact upon request.

## EXPERIMENTAL MODEL AND SUBJECT DETAILS

Five adult *Hipposideros armiger* (3 males, 2 females) weighing 60-70 g served as subjects for this study. The bats were caught with a hand net at a cave in Chongyang County, Xianning, Hubei Province, China in September 2022. Bats were housed in social groups of two to four individuals, with a regulated air temperature of around 24°C, relative humidity of about 60%, and a reversed light regime of 12 h of darkness and 12 h of light. Bats had free access to water and food at the home cage. All experimental procedures were approved by the Institutional Animal Care and Use Committee of the Central China Normal University.

## METHOD DETAILS

### Animal preparation and electrode array implant

For each of the five *H. armiger*, we implanted in the right inferior colliculus (IC) a 16-channel chronic recording silicon probe (A4×4-4mm-200-200-1250-H16_21mm, Neuronexus, USA, **Figure S1**) that was mounted on a modified nano-Drive (Cambridge NeuroTech, UK). The impedance of the silicon electrodes was 0.2∼0.6 MΩ. The IC of *H. armiger* sits on the dorsal surface of the brain, and all electrodes were implanted through the center of the bright spot that was histologically verified to be the central portion of the IC in another study (**Figure S1**). A few days before the implant surgery, we started to acclimate the bat to vocalize and to receive water and food by hanging freely at the perch within the recording chamber for about half an hour daily (i.e., habituation phase).

During the electrode implant surgery, the bat was deeply anesthetized using isoflurane mixed with oxygen (1.5% for induction; 0.5–1.2% for maintenance), with its head mounted on a stereotaxic apparatus (68810, RWD, China). A rostro-caudal incision was made to the region around the ICs and then the skins and muscles overlaying the ICs were pushed laterally to expose the skulls for electrode implants. A small elliptical craniotomy above the IC of the right hemisphere was performed using a dental drill (78001, RWD, China). After performing the craniotomy and durotomy, the moving parts of the nano-drive and the probe were covered with vaseline. The nano-drive was then slowly and carefully lowered to insert the tips of the electrodes into the IC until the bottom of the nano-drive gently touched the skull. The tips of the probe were initially positioned at a depth of approximately 1000 µm below the IC surface. The rest parts of the electrode assembly were then firmly glued to the skull with bone cement (C&B Metabond, Parkell, USA). The ground and reference wires were implanted into the left cerebral cortex and fixed with bone cement. The total weight of the surgical implants, including the silicon probe, nano-drive, and bone cement, was about 1.5 g. The rectal temperature of the bat was maintained at around 30°C through a regulated heating pad. The bat’s breathing rate was regularly noted every 15 minutes and was used to adjust the isoflurane level to ensure a proper level of anesthesia. Following surgery, erythromycin ointment was applied to the muscles/skin of the surgical site, and metacam (12.5 mg/kg) was administered orally for two consecutive days to prevent infection. The implanted bat was housed separately in a cage and allowed several days to recover before starting the neural recording experiment.

### Neural recording, sound recording, and playback

Neural signals and echolocation calls were recorded synchronously from freely vocalizing bats hanging from a perch hooked to the ceiling of a cage (0.6 m length × 0.4 m width × 1.2 m height) within an acoustic chamber (3 m length × 2 m width × 2.3 m height). The walls of the cage were constructed by vertically hanging velvet felt stripes (∼3.5 cm wide and 0.15 cm thick) spaced 15 cm apart. These stripes helped to reduce the likelihood of the bat flying away but only returned very weak echoes to the bat. The walls and ceiling were covered by 8-cm thick acoustic foam and the floor by a nylon blanket to reduce echoes and reverberations. Echolocation calls were recorded by an ¼ inch free-field measurement microphone (46BF-1, GRAS, Denmark) and were placed 20 cm away from the head of the freely hanging bat. When there were playback tasks, sound recording and playback were achieved via a multi-channel audio interface (Fireface 802, RME, Germany) at a sample rate of 192 kHz. Synchronization between the neural recording and sound recording and playback was achieved by regularly outputting a 1-ms long sine wave from one of the analog output channels every 0.1 seconds, which was recorded by a digital channel of the neural recording system. When there was no playback task, sound recording was sampled at a rate of 500 kHz with a multi-channel soundcard (PXIe-7858R, National Instruments, USA) with a customer-made LabVIEW program ^22^. Synchronization between the neural recording and sound recording was achieved by regularly outputting TTL signals from one of the analog output channels every 0.1 seconds, which were recorded by a digital channel of the neural recording system. A 1-inch sized electrostatic loudspeaker (ES1, TDT, USA) next to the microphone was used for sound playback. We constantly monitored the echolocation calls of the bat online with an ultrasound detector that converts the ultrasonic signals into audio sounds (D100, Pettersson Elektronik AB, Sweden). An infrared video camera (FDR-AX700, SONY, Japan), aided by an external IR light source (3022HW, Maixiu, China), was used to monitor the behavior of the bat, with the video signal displayed in real-time with a monitor outside the recording chamber.

The 16-channel neural signals were amplified and digitized via a tethered headstage at a sampling rate of 30 kHz, viewed online, and streamed to a computer (NeuroLego, BrainTech, Nanjing, China). At each electrode depth, we proceeded to record the neural signals when any electrode channel returned reasonably high-quality spikes. We recorded the neural responses of the putative neurons to spontaneously self-produced echolocation calls when the bat showed relatively high vocal activities. When the bat was in a relatively quiet state, we recorded the neural activities of the bat by playing either pure tones or pre-recorded echolocation calls. If none of the electrode channels showed spikes for a planned recording day, the electrodes were advanced ventrally by 50 µm, and no recording was made until the next recording day. During the recording, we paid particular attention to the neurons of relatively high signal-to-noise ratio (SNR) and made notes of their activities across the entire recording session. Information including whether a putative neuron changed the spike amplitude or was lost during the recording was noted down and used to guide further data analysis. While we observed frequent, yet small variations in the spike amplitude during the recording, a completely lost of a putative neuron occurred only before the fall of the implanted probe.

Sound playback, including both pure tones and the pre-recorded sonar vocalizations (vocalization playback, PLB), was controlled by a customer-made program in MATLAB (R2019a, Mathworks, USA), interfacing with a multiple-channel audio interface (Fireface 802, RME, Germany). Pure tones were generated online. The frequencies of pure tones ranged between 5 and 90 kHz. A frequency step of 0.1 kHz was used to generate pure tones for the frequencies between 33 and 36 kHz and between 68 and 72 kHz, which corresponded to the frequency areas of the 1^st^ and 2^nd^ CFs. For other frequency ranges, a step size of 1 kHz was used. The duration was 10 ms and the peak amplitude at the position of the bat was 75 dB SPL. Each pure tone was repeated 40 times with an inter-stimulus interval of 200 ms. Echolocation calls used for vocalization playback were prepared based on sound recordings from the same bat during the habituation phase before the implant surgery.

We built an ultrafast online acoustic playback setup based on the FPGA technique, which allowed us to broadcast a copy of the emitted vocalizations, as well as any pre-generated sound stimulus to the vocalizing bat at a controllable time delay down to ∼1 ms ^22^. We online broadcasted the recorded vocalizations at delays of 80 ms at one of three levels (approximately 40, 60, and 80 dB SPL). For offline playback (i.e. the vocalization playback condition), for each bat, a total of 10 call sequences were randomly selected and each call sequence contained 30 consecutive calls as well as a copy of the emitted vocalizations at 80 ms delay after each call. The vocalizations used for playback were band-pass filtered by 5-90 kHz, which contained both the 1^st^ and 2^nd^ harmonics. We standardized the call sequences by normalizing the amplitude of each call to one volt and the interval between consecutive calls to 0.12 seconds. In this study, only data from self-generated vocalization and offline playback conditions were included (**Figure S2**). Call sequences were broadcasted at a peak amplitude of 78 dB SPL at the location of the bat and each sequence was repeated four times. During sound playback, sound stimuli were amplified by a power amplifier (ED1, TDT, USA) before sending to the ultrasonic electrostatic speaker. The playback levels of both the pure tones and the vocalizations were probably comparable to or even slightly greater than the perceived amplitude of the self-produced calls. The source level of the self-produced calls (0.1 m in front of the bat) was 97 ± 9 dB and 111 ± 2 dB SPL, based on all recorded echolocation calls and the top 10% of the loudest calls from these bats. It was estimated that for echolocating bats the middle ear contraction and possible neural attenuation typically reduce the perceived call amplitude by about 30 - 40 dB during vocalization^43–45^. We achieve a flat frequency response for the playback system (± 1 dB) by digitally adjusting the output voltage of the playback stimuli. The procedures for calibrating and compensating the recording and playback systems were detailed in previous studies ^42,46^.

## QUANTIFICATION AND STATISTICAL ANALYSIS

### Data analysis

All data processing and analysis were performed using MATLAB (R2021a, MathWorks, USA). We performed offline spike sorting using the Wave_Clus3, a freely available toolbox from GitHub (https://github.com/ds7711/Vicario_lab_wave_clus3) ^47^. Spike sorting was performed for each type of acoustic stimuli independently, as we observed small, but visible variations of spike amplitude of some putative neurons during the recording. During spike sorting, the neural recordings were band-pass filtered between 600 and 3000 Hz (4^th^-order band-pass elliptic filter with 0.1 dB of passband ripple and 40 dB stopband attenuation). Single units were classified based on a minimum SNR of 14 dB and the presence of a refractory period. The sorted single units were all manually checked by Matlab’s toolbox MLIB6 to ensure quality (**Figure S3**).

A total of 106 single units from 5 bats were identified and included in the analysis. Each unit was clearly visible across the entire recording session that typically lasted for a few hours. Although our chronic preparation potentially allowed us to study the neuronal activity of the same neuron across days, the present experiment only studied and compared the neuronal activities within a single recording day. Temporal patterns of vocalization-related or playback-related activities were calculated as post-stimulus time histograms (PSTHs, 1 ms bin size) aligned by stimulus onset. Firing density refers to the average firing rate of one neuron over multiple calls based on the one-millisecond bin size. Significant changes from the baseline were estimated by rank-sum test with false discovery rate (FDR) correction. Inter-pulse-interval (IPI) of vocalizations greater than 120 ms were used for analyzing the neural responses to self-produced calls to avoid the effect of online playbacks on the response (**Figure S2, gray calls**). The best frequency (BF) was the frequency at which reached the maximum spike counts within 40 ms after the pure tone onset. Response peaks referred to the frequencies whose spike counts were more than three times the standard deviation above the background firing. The spike counts per stimulus were the average number of spikes within 40 ms of the onset of the vocalization or pure tone. Latency was calculated based on PSTH with a linear interpolation of 0.1 ms with over three times the standard deviation above the background firing. To compare the response latency, the maximal relative firing density, and the spike counts per stimulus of each IC neuron between vocalization and vocalization playback conditions, for each neuron we randomly resampled 100 stimuli, 20 stimuli for pure tone playback condition, from the entire corresponding datasets and repeated it 30 times, and then evaluated the statistical significance. We performed the principal component analysis (PCA) using the spike data (40 ms PSTH after the onset of vocalizations or vocalization playbacks). We used the normalized cross-correlation method to compare similarity in firing patterns. Specifically, we performed the similarity analysis with 50% of the vocalizations and 50% of the playback vocalizations randomly sampled from the entire experimental datasets. Then we calculated the maximum correlation coefficient for the firing patterns between the resampled 50% vocalization dataset and original 100% vocalization dataset (within-group), between the resampled 50% playback vocalization dataset and the original 100% playback dataset (within-group), and between the resampled 50% vocalization dataset and the resampled 50% playback vocalization dataset (between-group). A coefficient value closer to 1 indicates a more similar firing pattern. All statistical tests were performed using the non-parametric Wilcoxon rank-sum test unless otherwise stated. Correlation analysis was based on the Pearson correlation. The significance level was set to 0.05.

---

**Figure S1.**
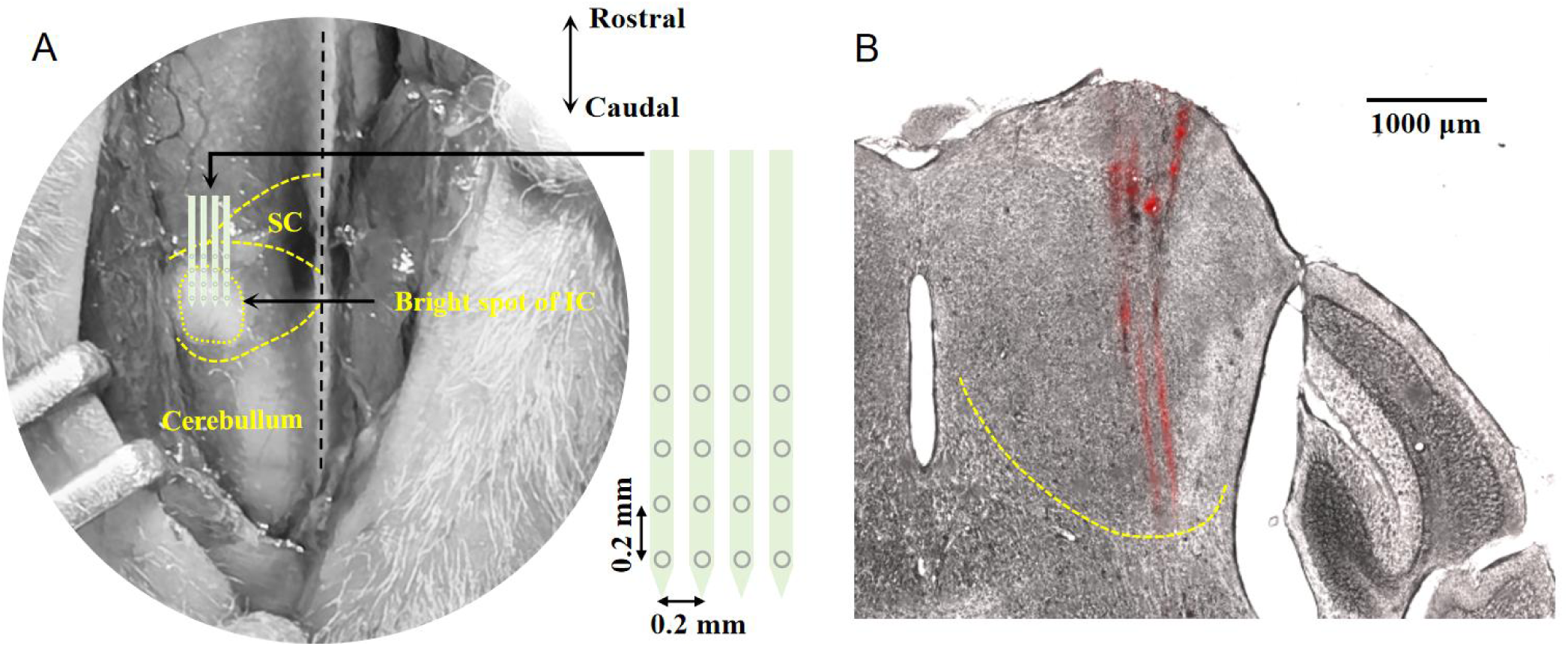
Top view of the skull surface and histological verification of the central inferior colliculus (ICc) in *Hipposideros armiger*. (A) The IC of *H. armiger* sits on the dorsal surface of the brain and manifests as a bright spot. All silicon probes (4 × 4 configuration) were implanted through the center of the bright spot (left panel). SC, superior colliculus. (B) The center of the bright spot in the skull surface corresponds to the ICc. Tracks of a 4 × 4 electrode array inserted via the center of the bright spot, as in the case of the implanted silicon probes. The electrode array was coated with fluorescent Dil dye before the penetration.

**Figure S2.**
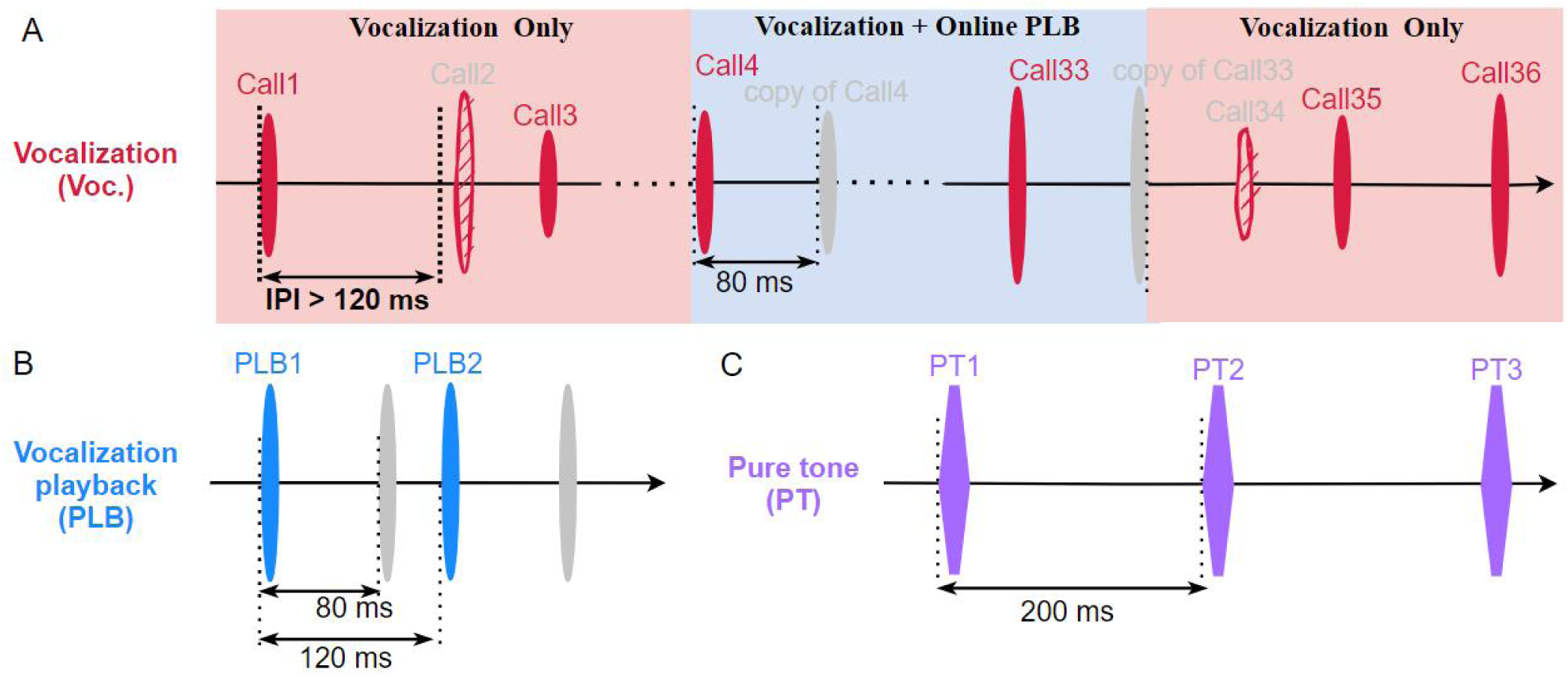
Schematic illustration of the three acoustic stimulation conditions. **(A)** During the vocalization condition (Voc.), the bat spontaneously produced echolocation calls (red). The vocalization condition includes an online playback (Voc. + Online PLB, blue background filled) window preceded and succeeded by a 2-second vocalization window without playback (Voc. Only, pink background filled). The online playback window contained a total of 30 calls and each call was followed by a playback (copy) of the call at a delay of 80 ms. As the neural responses rarely persisted for 40 ms relative to the onset of the stimulus, we used 120 ms as the minimum inter-call interval for analyzing the neural responses of self-produced vocalizations. In this illustration, Call2 and Call34 represent excluded calls. (B) The vocalization playback condition (PLB) matched the vocalization condition using a sequence of the recorded echolocation calls (blue), and each echolocation call was followed by a copy of the attenuated call (grey). The inter-call interval was fixed to 120 ms. (C) The pure tone condition included sequences of pure tones at an interval of 200 ms. Both vocalization playback and pure tone tasks were performed when the bat was not actively vocalizing.

**Figure S3.**
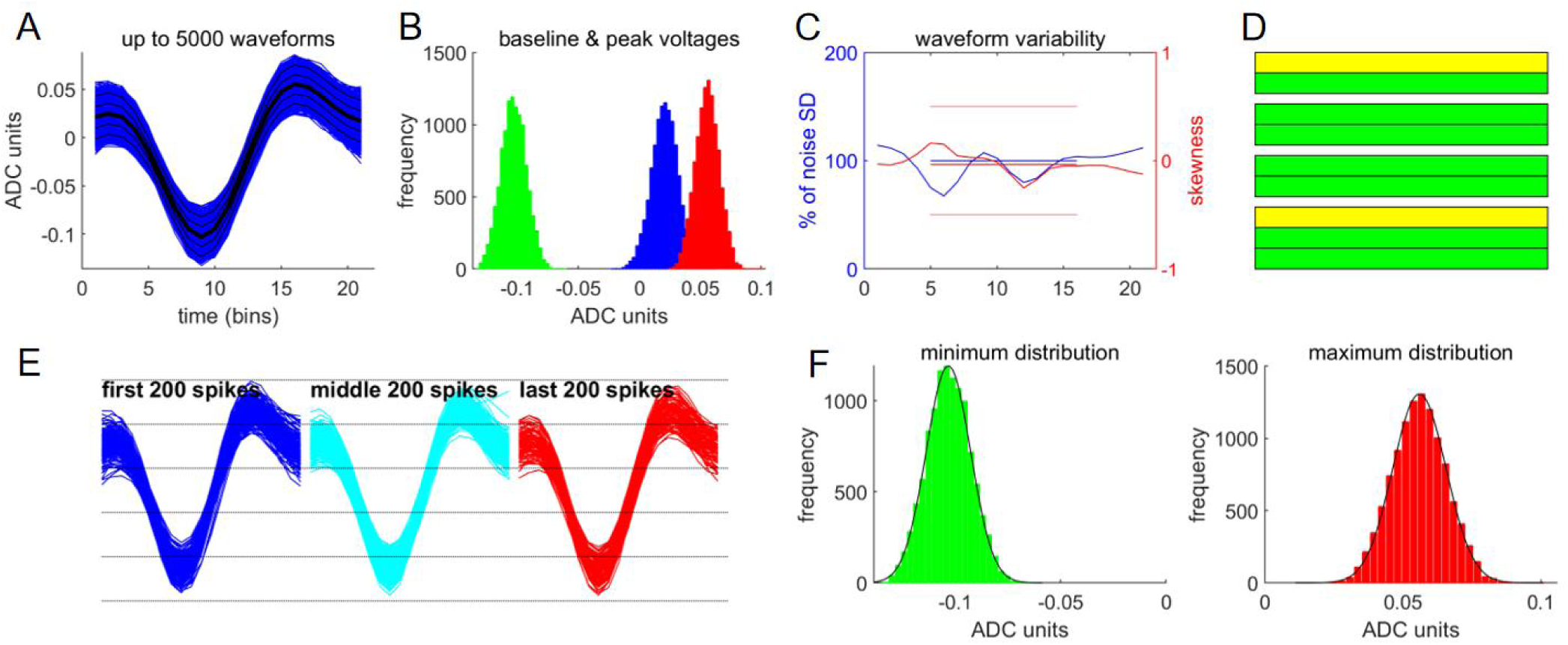
Quality verification of isolated single units using Matlab’s toolbox MLIB6. (A) Individual and average waveform of an isolated single unit. (B) The frequency distributions of the noise amplitude (first bin of all waveforms, blue), the negative peak amplitudes (green), and the positive peak amplitudes (red). Generally, all distributions of single units should be approximately normal and separated from each other. (C) The SD (blue) and the skewness (red) of the voltage distributions as a function of time (waveform ticks). Solid lines denote mean SD and mean skewness. The thin red line was the skewness of ± 0.5. Ideally, skewness should be 0, but variation within these boundaries was deemed acceptable for single neurons’ waveform variability, and standard deviation was expressed as percentages of the mean noise SD (SD in the first five ticks), and the lower its variability, the better. (D) Unit quality SNR is reflected by the colored traffic lights, with green denoting the optimal or high-quality, yellow for worth analyzing, and red for low-quality single units. (E) Waveforms of the first, middle, and last 200 spikes of the recording file. (F) The distribution of the minimum amplitudes and maximum amplitudes of all spike waveforms and the fitted curves by normal distribution.

**Tabel S1.**
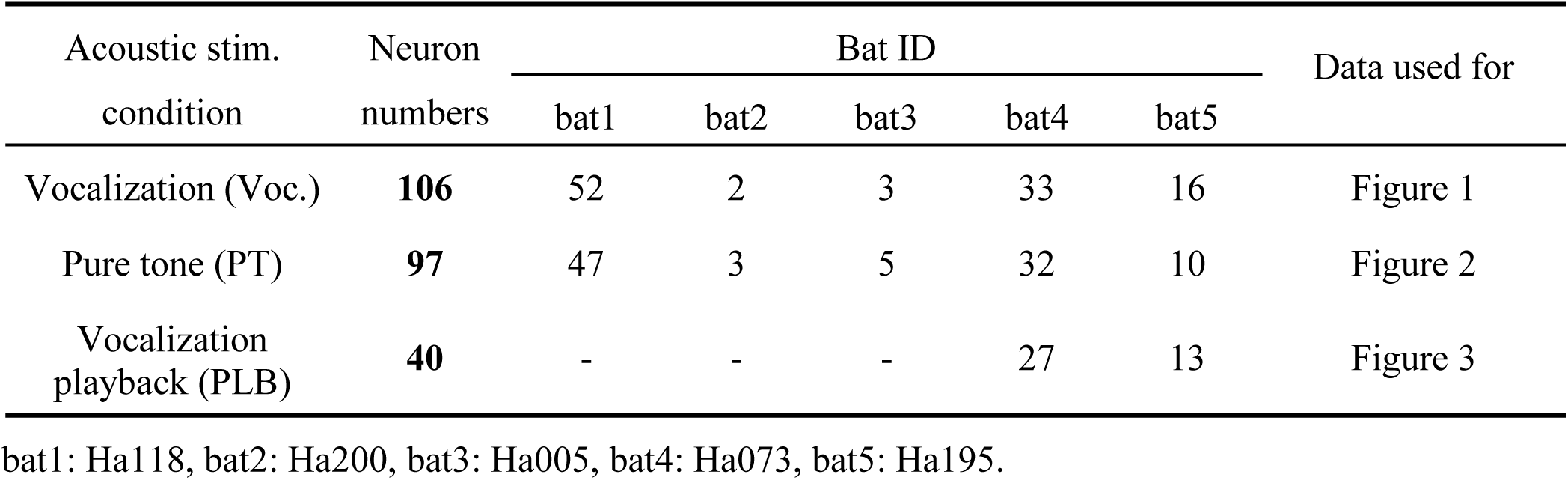
Details for the 106 inferior colliculus neurons from five implanted bats.

**Movie S1 An example inferior colliculus neuron reversed the response polarity, from no excitatory response to self-produced vocalizations to vocalization-playback-induced excitation.** The top left panel shows the bandpass-filtered neural recording for spikes (600 – 3,000 Hz). The bottom left panel shows the spectrogram (5 – 90 kHz filter) of the self-produced echolocation calls in the vocalization condition or the vocalization playbacks. The top right panel displays the waveforms of the superimposed spikes for the isolated single unit. The bottom right panel displays the vocalization-onset-aligned spikes (raster plot). Note, that this example neuron (Ha073_04r62_chn15, 2610 μm recording depth) showed spontaneous firing that was not related to the vocalization or vocalization playback, which was typical for the sampled IC neurons during the behavioral/awake setup. Both the audio of the echolocation calls and the neural recordings were slowed down by a factor of 16 times.

## References

1. Schneider, D.M., and Mooney, R. (2018). How movement modulates hearing. Annu Rev Neurosci 41, 553–572.

2. Eliades, S.J., and Wang, X. (2019). Corollary discharge mechanisms during vocal production in marmoset monkeys. Biol. Psychiatry Cogn. Neurosci. Neuroimaging 4, 805–812.

3. Brainard, M.S., and Doupe, A.J. (2000). Auditory feedback in learning and maintenance of vocal behaviour. Nature Reviews Neuroscience 1, 31–40.

4. Nieder, A., and Mooney, R. (2019). The neurobiology of innate, volitional and learned vocalizations in mammals and birds. Philosophical Transactions of the Royal Society B: Biological Sciences 375, 20190054.

5. Eliades, S.J., and Wang, X. (2003). Sensory-motor interaction in the primate auditory cortex during self-initiated vocalizations. Journal of Neurophysiology 89, 2194–2207.

6. Eliades, S.J., and Wang, X. (2008). Neural substrates of vocalization feedback monitoring in primate auditory cortex. Nature 453, 1102–1106.

7. Eliades, S.J., and Tsunada, J. (2018). Auditory cortical activity drives feedback-dependent vocal control in marmosets. Nat Commun 9, 2540.

8. Tsunada, J., Wang, X., and Eliades, S.J. (2024). Multiple processes of vocal sensory-motor interaction in primate auditory cortex. Nat Commun 15, 3093.

9. Aliu, S.O., Houde, J.F., and Nagarajan, S.S. (2009). Motor-induced Suppression of the Auditory Cortex. Journal of Cognitive Neuroscience 21, 791–802.

10. Chang, E.F., Niziolek, C.A., Knight, R.T., Nagarajan, S.S., and Houde, J.F. (2013). Human cortical sensorimotor network underlying feedback control of vocal pitch. Proceedings of the National Academy of Sciences 110, 2653–2658.

11. Rummell, B.P., Klee, J.L., and Sigurdsson, T. (2016). Attenuation of responses to self-generated sounds in auditory cortical neurons. Journal of Neuroscience 36, 12010–12026.

12. Schneider, D.M., Sundararajan, J., and Mooney, R. (2018). A cortical filter that learns to suppress the acoustic consequences of movement. Nature 561, 391–395.

13. Winer, J.A., and Schreiner, C.E. eds. (2005). The Inferior Colliculus (Springer).

14. Gruters, K.G., and Groh, J.M. (2012). Sounds and beyond: multisensory and other non-auditory signals in the inferior colliculus. Front. Neural Circuits 6, 96.

15. Olthof, B.M.J., Rees, A., and Gartside, S.E. (2019). Multiple nonauditory cortical regions innervate the auditory midbrain. J. Neurosci. 39, 8916–8928.

16. Gartside, S.E., Rees, A., and Olthof, B.M. (2023). Motor, somatosensory, and executive cortical areas directly modulate firing activity in the auditory midbrain. Preprint at bioRxiv.

17. Tammer, R., Ehrenreich, L., and Jürgens, U. (2004). Telemetrically recorded neuronal activity in the inferior colliculus and bordering tegmentum during vocal communication in squirrel monkeys (Saimiri sciureus). Behavioural Brain Research 151, 331–336.

18. Schuller, G. (1979). Vocalization influences auditory processing in collicular neurons of the CF-FM-bat, *Rhinolophus ferrumequinum*. J Comp Physiol A 132, 39–46.

19. Schoeppler, D., Schnitzler, H.-U., and Denzinger, A. (2018). Precise Doppler shift compensation in the hipposiderid bat, *Hipposideros armiger*. Sci. Rep. 8, 4598.

20. Zhang, Y., Lin, A., Ding, J., Yang, X., Jiang, T., Liu, Y., and Feng, J. (2019). Performance of Doppler shift compensation in bats varies with species rather than with environmental clutter. Anim. Behav. 158, 109–120.

21. Lu, M., Zhang, G., and Luo, J. (2020). Echolocating bats exhibit differential amplitude compensation for noise interference at a sub-call level. J. Exp. Biol. 223, 225284.

22. Wang, H., Zhou, Y., Li, H., Moss, C.F., Li, X., and Luo, J. (2022). Sensory error drives fine motor adjustment. Proc Natl Acad Sci USA 119, e2201275119.

23. Sutter, M.L., and Schreiner, C.E. (1991). Physiology and topography of neurons with multipeaked tuning curves in cat primary auditory cortex. Journal of Neurophysiology 65, 1207–1226.

24. Kadia, S.C., and Wang, X. (2003). Spectral integration in A1 of awake primates: Neurons with single- and multipeaked tuning characteristics. Journal of Neurophysiology 89, 1603–1622.

25. Manley, J.A., and Müller-Preuss, P. (1981). A Comparison of the Responses Evoked by Artificial Stimuli and Vocalizations in the Inferior Colliculus of Squirrel Monkeys. In Neuronal Mechanisms of Hearing, J. Syka and L. Aitkin, eds. (Springer US), pp. 307–310.

26. Casseday, J., Fremouw, T., and Covey, E. (2002). The inferior colliculus: A hub for the central auditory system. In Integrative Functions in the Mammalian Auditory Pathway Springer Handbook of Auditory Research., D. Oertel, R. Fay, and A. Popper, eds. (Springer New York), pp. 238–318.

27. Suga, N. (2015). Neural processing of auditory signals in the time domain: Delay-tuned coincidence detectors in the mustached bat. Hear Res 324, 19–36.

28. Wohlgemuth, M.J., Luo, J., and Moss, C.F. (2016). Three-dimensional auditory localization in the echolocating bat. Curr Opin Neurobiol 41, 78–86.

29. Macias, S., and Llano, D.A. (2023). Descending projections to the auditory midbrain: evolutionary considerations. J Comp Physiol A 209, 131–143.

30. Geberl, C., Brinkløv, S., Wiegrebe, L., and Surlykke, A. (2015). Fast sensory–motor reactions in echolocating bats to sudden changes during the final buzz and prey intercept. Proc. Natl. Acad. Sci. U.S.A. 112, 4122–4127.

31. Luo, J., and Moss, C.F. (2017). Echolocating bats rely on audiovocal feedback to adapt sonar signal design. Proc Natl Acad Sci USA 114, 10978–10983.

32. Luo, J., Kothari, N.B., and Moss, C.F. (2017). Sensorimotor integration on a rapid time scale. Proc. Natl. Acad. Sci. USA 114, 6605–6610.

33. Pedersen, M.B., Egenhardt, M., Beedholm, K., Skalshøi, M.R., Uebel, A.S., Hubancheva, A., Koseva, K., Moss, C.F., Luo, J., Stidsholt, L., et al. (2024). Superfast Lombard response in free-flying, echolocating bats. Current Biology 34, 2509–2516.e3.

34. Yang, Y., Lee, J., and Kim, G. (2020). Integration of locomotion and auditory signals in the mouse inferior colliculus. eLife 9, e52228.

35. Schneider, D.M., Nelson, A., and Mooney, R. (2014). A synaptic and circuit basis for corollary discharge in the auditory cortex. Nature 513, 189–194.

36. Audette, N.J., Zhou, W., La Chioma, A., and Schneider, D.M. (2022). Precise movement-based predictions in the mouse auditory cortex. Current Biology 32, 4925–4940.e6.

37. Müller-Preuss, P. (1983). Inhibition of auditory neurons during phonation: Evidence of feed-forward mechanisms in brain processes controlling audio-vocal behavior? In Advances in Vertebrate Neuroethology NATO Advanced Science Institutes Series., J.-P. Ewert, R. Capranica, and D. Ingle, eds. (Springer US), pp. 919–923.

38. Slee, S.J., and David, S.V. (2015). Rapid task-related plasticity of spectrotemporal receptive fields in the auditory midbrain. J. Neurosci. 35, 13090–13102.

39. De Franceschi, G., and Barkat, T.R. (2021). Task-induced modulations of neuronal activity along the auditory pathway. Cell Reports 37, 110115.

40. Salles, A., Loscalzo, E., Montoya, J., Mendoza, R., Boergens, K.M., and Moss, C.F. (2024). Auditory Processing of Communication Calls in Interacting Bats. iScience, 109872.

41. Wetekam, J., Hechavarría, J., López-Jury, L., González-Palomares, E., and Kössl, M. (2024). Deviance detection to natural stimuli in population responses of the brainstem of bats. J Neurosci 44, e1588232023.

42. Ye, H., and Luo, J. (2022). Perceptual hearing sensitivity during vocal production. iScience 25, 105435.

43. Henson, O.W. (1965). The activity and function of the middle-ear muscles in echo-locating bats. The Journal of Physiology 180, 871–887.

44. Suga, N., and Jen, P.H. (1975). Peripheral control of acoustic signals in the auditory system of echolocating bats. Journal of Experimental Biology 62, 277–311.

45. Suga, N., and Schlegel, P. (1972). Neural attenuation of responses to emitted sounds in echolocating bats. Science 177, 82–84.

46. Ma, N., Xia, H., Yu, C., Yin, K., and Luo, J. (2024). Effects of insect pursuit on the Doppler shift compensation in a hipposiderid bat. J Exp Biol 227, jeb246355.

47. Quiroga, R.Q., Nadasdy, Z., and Ben-Shaul, Y. (2004). Unsupervised spike detection and sorting with wavelets and superparamagnetic clustering. Neural Comput 16, 1661–1687.

